# Extracellular matrix stiffness regulates force transmission pathways in multicellular ensembles of human airway smooth muscle cells

**DOI:** 10.1101/402842

**Authors:** Samuel R. Polio, Suzanne E Stasiak, Ramaswamy Krishnan, Harikrishnan Parameswaran

**Affiliations:** Dept. of Bioengineering, Northeastern University, Boston MA 021115; Beth Israel Deaconess Medical Center, Harvard Medical School, Boston MA 021115

## Abstract

For an airway or a blood vessel to narrow, there must be a connected path that links the smooth muscle (SM) cells with each other, and transmits forces around the organ, causing it to constrict. Currently, we know very little about the mechanisms that regulate force transmission pathways in a multicellular SM ensemble. Here, we used extracellular matrix (ECM) micropatterning to study force transmission in a two-cell ensemble of SM cells. Using the two-SM cell ensemble, we demonstrate (a) that ECM stiffness acts as a switch that regulates whether SM force is transmitted through the ECM or through cell-cell connections. (b) Fluorescent imaging for adherens junctions and focal adhesions show the progressive loss of cell-cell borders and the appearance of focal adhesions with the increase in ECM stiffness (confirming our mechanical measurements). (c) At the same ECM stiffness, we show that the presence of a cell-cell border substantially decreases the overall contractility of the SM cell ensemble. Our results demonstrate that connectivity among SM cells is a critical factor to consider in the development of diseases such as asthma and hypertension.

## Introduction

Asthma, Crohn’s disease, and hypertension are widespread chronic diseases that affect the airways, the gut, and blood vessels respectively. Although they are very different diseases, physiologically they share one common feature: they are all characterized by excessive narrowing of the luminal area of a hollow, tubular organ. Traditionally, these changes have been viewed as a disease of the smooth muscle (SM) and treatment strategies have focused on identifying and targeting molecular pathways that control force generation inside the SM cell^1–4^. However, it is becoming increasingly clear that the etiology of excessive narrowing of airways and blood vessels cannot be explained by considering the SM cell in isolation from its native environment^5–9^. In order to generate enough force to effect a change in the luminal area, the contractile apparatus of individual SM cells must physically connect with each other to form a force transmission pathway that wraps around the circumference of the tube. Clearly, the number of SM cells that form this connected path as well as the stiffness of the cell-cell and cell-extracellular matrix interactions will dictate how much luminal constriction is achieved for a given level of SM activation. Currently, we know very little about how these connections among SM cells are achieved in a multicellular SM ensemble.

At the cellular level, SM cells can form direct connections with their neighboring cells through cadherin-mediated adherens junctions^10^. They can also connect to the extracellular matrix (ECM) through integrin-mediated focal adhesions^11^. Focal adhesions allow SM cells to indirectly connect with other SM cells over long distances through the ECM^12,13^. As such, SM cells *in vivo* have multiple options available to them to make connections among themselves and transmit their force. The factors that dictate the choice of force transmission pathways used by SM cells in healthy and diseased tissue is still unclear. Both focal adhesions and adherens junctions are mechanosensitive structures through which cells can respond to, and probe the stiffness and ligands present in their surrounding environment. Focal adhesions size and maturation rates have been shown to depend on cytoskeletal tension^14^ and ECM stiffness^15^. Similarly, in cell-cell cadherin junctions, the cadherin-catenin complex/actin filament binding in adherens junctions has been shown to exhibit catch bond characteristics up to 10pN after which it transitions into a slip bond^16^. Based on these data, we hypothesize that mechanical cues such as ECM stiffness can alter the nature of force transmission pathways (cell-cell vs cell-ECM) in a multicellular ensemble of human SM cells.

To test this hypothesis, we applied ECM micropatterning techniques to create islands of two human airway smooth muscle (ASM) cells and measured the effect of changing ECM stiffness on the ASM force transmitted through cell-cell coupling^17,18^. We report direct measurement of forces exerted by an ASM cell on its neighbor, and on the ECM for substrates with stiffness matching healthy (Young’s modulus, E=300Pa) and remodeled tissue (Young’s modulus, E=13 kPa). On substrates with the stiffness matching that of healthy tissue, we find that ASM cells exert more of their longitudinal tension on their neighboring ASM cells compared to the ECM. On soft substrates, imaging reveals the presence of well defined adherens junctions connecting ASM cells indicating that there is strong coupling between the cells in healthy tissue. As the substrate stiffness is increased to match that of remodeled tissue, ASM-ASM coupling weakens and more of the ASM force is exerted on the matrix. Imaging confirms the gradual loss of adherens junctions and replacement by focal adhesions as the ECM stiffens. These experiments indicate that the ECM stiffness can act as a switch that regulates whether forces are transmitted via the ECM or through cell-cell contacts. The change in connectivity can also significantly change the overall contractile strength of the ensemble. Excessive contraction of airways and bloodvesels can therefore emerge as a result of change in connectivity among SM cells driven by extracellular matrix remodeling. Our results highlight the need to develop new therapies for asthma and hypertension that target extracellular matrix remodeling.

## Results

### Creating a two-cell ensemble of human airway smooth muscle cells

In order to measure the forces that SM cells exert on their neighbour and on the ECM, we adapted an experimental system that has been previously described for similar measurements in cardiac myocytes^17^. Briefly, the method involves creating a rectangular shaped “micro tissue” with exactly two cells in contact with each other. In the case of ASM cells, we wanted the cells to be elongated and aligned in a manner that was consistent with how ASM cells are organized *in vivo*^19^. To this end, we used a novel UV activated ECM patterning technique (Primo, Alvéole, France) to create rectangular shaped islands of gelatin on a dimethylpolysiloxane (PDMS) substrate^20^. To facilitate measurement of cellular traction forces, a layer of fluorescent beads was spin-coated on the surface of the gel substrate prior to gelatin patterning. The PDMS substrate material used in this study was NuSil^20^, an optically clear, non-porous material whose stiffness can be tuned in the range Young’s modulus, E =300Pa-40kPa. A substrate stiffness of E=300Pa was used to mimic the ECM stiffness of healthy human airways. With the onset of airway remodeling, there is collagen deposition in the airways and the stiffness of the ECM increases^21^. A substrate stiffness of E=13kPa was used to mimic remodeled ECM. The length of the rectangular pattern of gelatin was specified to match the length of ASM cells plated on non-patterned NuSil gels (150 µm) and the breadth (15 µm) was set by trial and error to allow for enough space within the rectangle to fit exactly two ASM cells. Fig 1A shows a composite image with the phase contrast image of a two-ASM cell ensemble superimposed on fluorescent images of their nuclei stained in blue using DAPI and the 150 µm x15 µm rectangular micro-pattern of the Nusil gel surface using fluorescent gelatin (green). A phase contrast image of a two ASM cell ensemble with displacement fields generated from the fluorescent beads overlaid on top is shown in Fig. 1B. We found that these dimensions (150 µm x15 µm) gave the cells enough area to spread and form a well-defined border between them. A detailed description of the protocols used for cell culture, gel manufacture and the ECM micro-patterning can be found in the Methods section.

**Figure 1.**
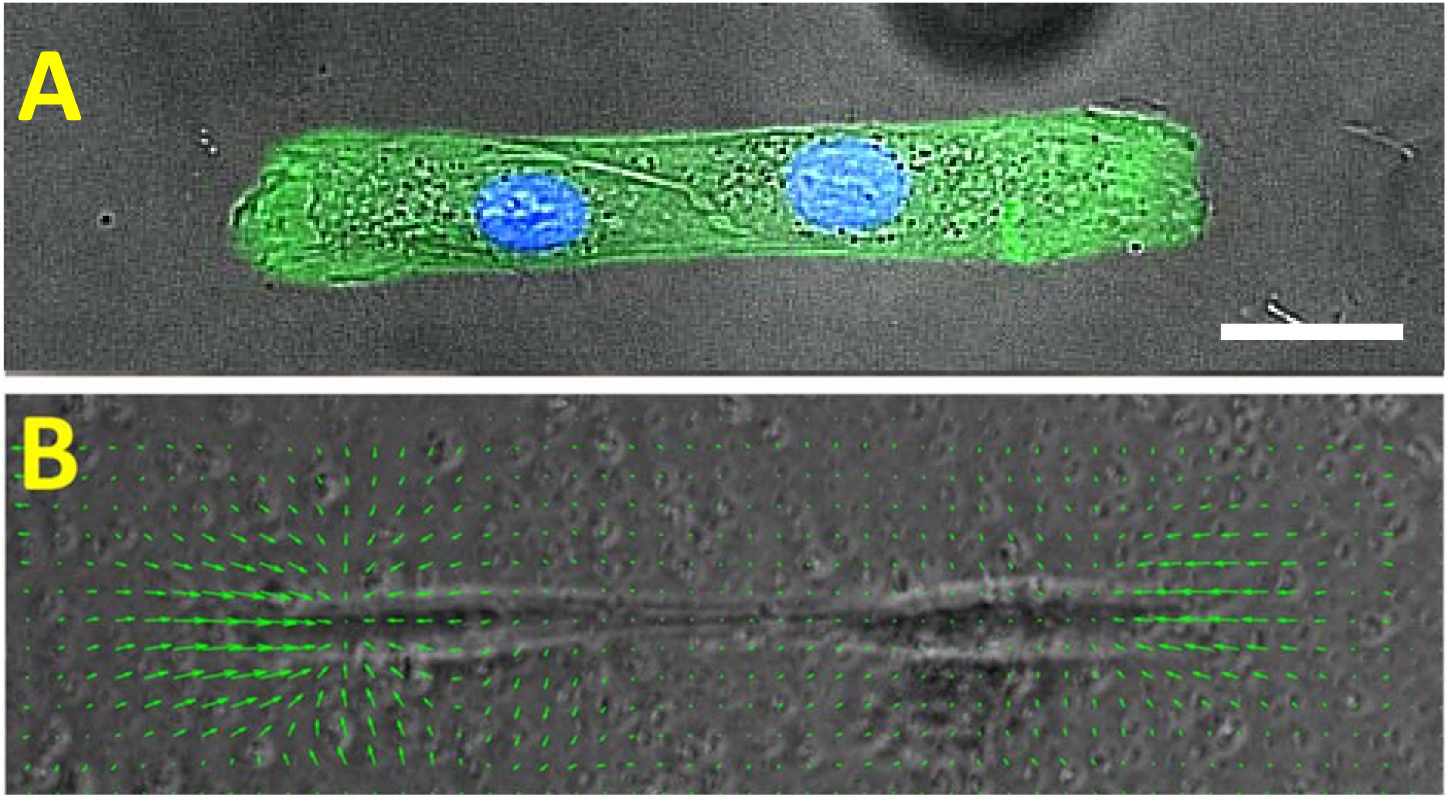
**(A)** shows a composite image with a phase contrast image of a two-ASM cell ensemble superimposed on fluorescent images of their nuclei stained in blue using DAPI. The 150 µm x15 µm rectangular micro-pattern on the Nusil gel surface is marked using fluorescent gelatin (green). The scale bar is 25µm. **(B)** shows the displacement field (green arrows) calculated from the movement of 1µm beads spin coated on the gel surface. The displacement field is superimposed on the phase contrast image of the two-SM cell pair.

### ASM-ASM coupling strength decreases with increasing ECM stiffness

Using the beads embedded in the NuSil gel, we measured traction forces exerted by the two-cell ensemble of ASM cells on the substrate. To measure the forces exerted by each ASM cell on its neighbor, we manually outlined the cell-cell junction from phase contrast images and used this to separate the two-ASM cell ensemble into two distinct regions corresponding to the each ASM cell. (Fig. 2A). This allowed us to directly calculate the force exerted by each cell on the cell-cell junction *F⃗_j cell_* as the unbalanced traction force in each cell^18,22^.

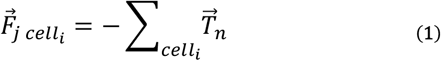

where *F⃗_j celli_* is the force exerted by the *i^th^* cell (*i=1,2*) on the cell-cell boundary. *T⃗_n_* is the traction force vector corresponding to computational grid *n* within the *i^th^* cell. The average magnitude of the force exerted on the cell-cell junction, 〈‖*F⃗_j cell_1__, F⃗_j cell_2__*‖〉 is plotted in Fig. 2C for healthy (E =300 Pa) and remodeled ECM (E =13kPa). We found that as the ECM stiffness increases, the force on the cell-cell junction increased significantly from 0.022±0.009 µN (N =6) to 0.179±0.073 µN (N =4) (p<0.05, Mann-Whitney). In Fig. 2B we show changes in the average longitudinal tension of the ASM cells, Nss as defined by

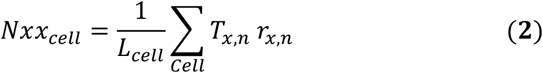

In our two-cell pairs at E =300 Pa and 13kPa, 〈‖*N_xx,cell_* ‖〉 increases significantly from 0.05±0.03µN (N =6) at 300 Pa to 0.48±0.2 µN (N =4) at 13kPa (p<0.05, Mann-Whitney). Several previous studies have noted that cellular traction forces increase with the increase in ECM stiffness^23,24^. Fig 2B confirms that within the range of ECM stiffness that is of interest to this study, changing ECM stiffness increases ASM cellular tractions significantly. Consequently, the force exerted on the cell-cell junction 〈‖*F⃗_j cell_1__, F⃗_j cell_2__* ‖〉 is not a reliable indicator of cell-cell coupling strength, as this quantity will always increase as traction forces ‖*T⃗_n_*‖ increase. To normalize for this effect, we calculate a dimensionless ratio (Ψ) which is indicative of the mechanical coupling between cells^17^ and is defined as:

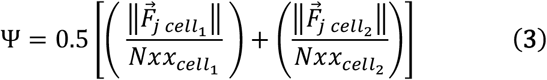

In these calculations, the *x*-axis was assumed to be aligned with the long axis of the rectangular pattern with the origin at the cell-cell border such that *r_x_* = 0 at the cell-cell junction. The length of the cell *L_cell_* was measured manually from phase contrast images as the length parallel to the *x-axis* (the direction in which the ASM cells are aligned). From equation (3), the ASM-ASM coupling strength, Ψ is a dimensionless ratio which measures the fraction of the cell’s longitudinal tension that it exerts on the boundary. For ASM cells, Ψ decreased significantly from 0.51±0.18 (N =6) to 0.28±0.05 (N =4) (p<0.05, t-test) as substrate stiffness was increased from 300Pa to 13kPa (Fig. 2D) indicating that the ASM cells can potentially physically decouple from each other as ECM stiffness is increased. To verify this result, we imaged the cells after fluorescently staining for markers of cell-cell and cell-ECM adhesion.

**Figure 2.**
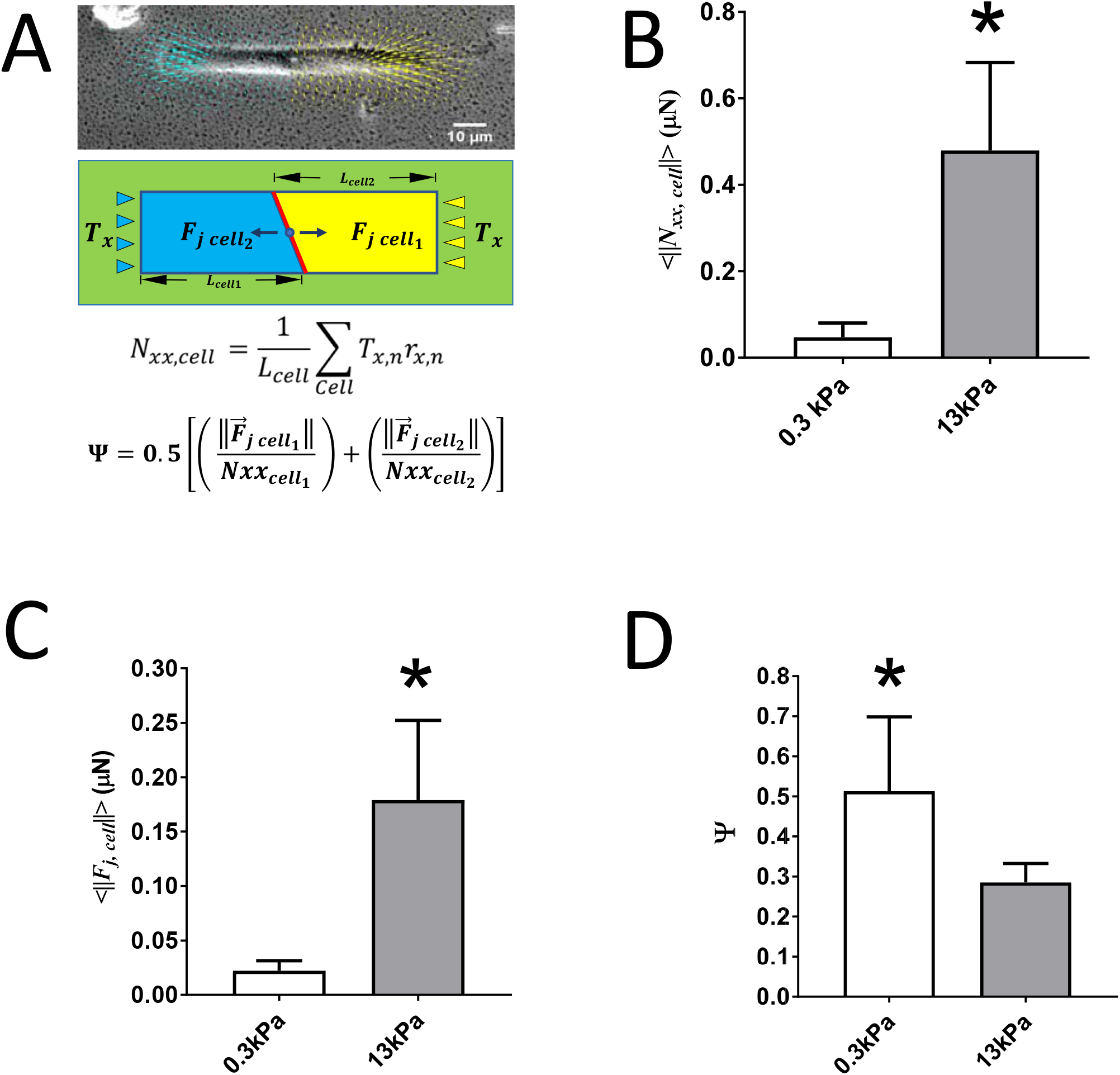
**(A)**The phase contrast image shows a two ASM cell pair, the traction field generated by this “micro-tissue” is split in two (yellow for cell1 and blue for cell2) by manually outlining each cell in the image. The force exerted on the cell-cell junction but each cell *F⃗_j cell_* can be calculated as the unbalanced traction force in that cell. **(B)** The average magnitude ofthe longitudinal tension of ASM cells in our two-cell ensemble〈‖*N_xx,cell_*‖〉(p<0.05; Mann-Whitney). **(C)** Average magnitude of the force exerted at the cell-cell junction 〈‖*F⃗_j,cell_* ‖〉 (p<0.05; Mann-Whitney). **(D)** Strength of the coupling between the two ASM cells, Ψ calculated using equation (3) (p<0.05; t-test).

### With the onset of remodeling cell-cell adherens junctions are progressively replaced by cell-ECM focal adhesions

To verify our mechanical measurements of cell-cell coupling strength, we imaged our two-ASM cell ensembles after staining for β-catenin (an adherens junction marker), vinculin (focal adhesion marker), and the nucleus (DAPI). Images of cells stained at three different stiffnesses are shown in Fig. 3. On substrates mimicking ECM stiffness of healthy tissue E =300Pa (Fig. 3A), the border between ASM cells is well defined by the β-catenin stain indicating the presence of adherens junctions at the ASM-ASM cell border. As ECM stiffness increases to E=13kPa, well defined focal adhesions (green: Fig. 3B) start to develop at the ASM-ASM cell border (yellow arrows in Panel B). This structural change is reflected in our mechanical measurements of Ψ which indicate a significant decrease in cell-cell coupling strength (Fig. 2). At E =40kPa, the adherens junctions and the cell-cell border completely disappear, and focal adhesions can be seen at the border (yellow arrows Fig. 3C). Now, the two ASM cell ensemble separates into two individual ASM cells that are connected to the ECM. There is no functional border (of adherens junctions) between the ASM cells at E=40kPa. It should be noted that vinculin can also appear in adherens junctions, but if it does, it would spatially co-localize with β-catenin—which is not the case here. Hence, we can be certain that the vinculin (green) in Figs. 3A-C represents focal adhesions.

**Figure 3.**
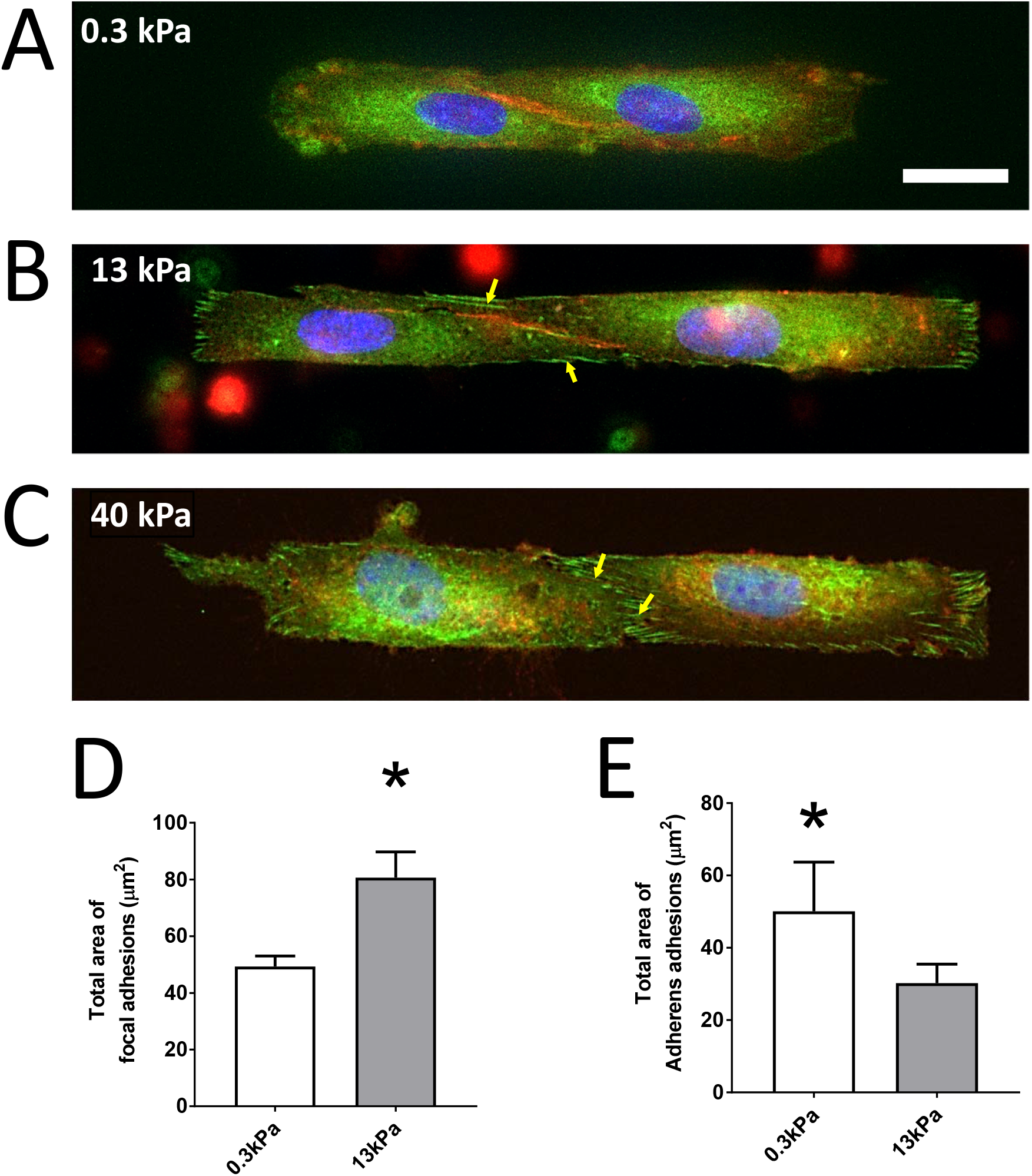
**(A-C)** ASM cells were stained for β-catenin (an adherens junction marker; red), Vinculin (focal adhesion marker; green), and the nucleus (DAPI; blue). Scale bar is 10µm. **(A) Healthy ECM, E =300Pa:** ASM-cells form stable cell-cell junctions and the boundary is clearly marked by β-catenin stain. **(B, C) Remodeled, stiffer ECM (E =13 kPa-40 kPa):** As the ECM stiffness increases to 13kPa, one sees the appearance of focal adhesions near the cell-cell boundary (marked by yellow arrows). When ECM stiffness is further increased to 40 kPa, the ASM-ASM boundary (red) gives way to cell ECM focal adhesions (green). **(D)** Total area occupied by focal adhesions (green) in two-cell pair increases significantly when ECM stiffness increases from 0.3 kPa to 13 kPa (p<0.05, t-test). **(E)** Total area occupied by adherens adhesions (red) in two-cell pairs decreases significantly when ECM stiffness increases from 0.3 kPa to 13kPa (p<0.05, t-test).

To test whether the loss of adherens junctions between 0.3kPa and 13kPa was significant, we selected the fluorescence channels for focal adhesions (green) and adherens junctions (red). These images were first thresholded based on grayscale intensity to identify focal adhesions and adherens junctions. To avoid counting regions produced by background fluorescence and noise, we manually masked out such artifacts. The total area of focal adhesions and adherens junctions were then measured by counting the number of pixels and then converting the area in pixels to units of µm^2^ using the appropriate scaling factors. Figs 3D and 3E show the change in total area of focal adhesions and adherens junctions respectively. There was a statistically significant increase in focal adhesion area as E was increased from 0.3kPa to 13kPa from 49.3 ± 3.7 µm^2^(N =5) to 80.7 ± 9.1 µm^2^(N =5) (p<0.05, t-test). Concurrently, the cell-cell adherens junction area underwent a statistically significant decrease in size from 50.1 ± 13.6 µm^2^(N =4) to 30.2 ± 5.2 µm^2^(N =4) (p<0.05, t-test). These quantitative measurements correspond to qualitative visual observations from fluorescent images (Figs 3A&B). Images at E=40kPa are not included in this comparison because at E =40kPa, there were no discernable adherens junctions that could be clearly identified by thresholding. Additionally, the bead displacements at E=40kPa were too low for reliable measurement of cellular traction forces.

### Clarifying the confounding effect of ECM stiffness on the interpretation of data

Our results thus far give a strong indication that in a remodeled extracellular environment, the ASM cells change their connectivity from ASM-ASM connections to ASM-ECM connections and the net force of the ASM ensemble increases significantly. However, there is considerable evidence that changes in ECM stiffness can also influence cell mechanics even in single cells^25–27^ and so cell-ECM interactions can be a potential confounding factor in interpreting our experimental findings. To isolate the effect of ASM-ASM coupling from ASM-ECM coupling on our results, we needed an independent means to verify that, *at the same ECM stiffness*, an ensemble of ASM cells that form strong adherens junctions among themselves would behave differently from an ensemble of adjoined ASM cells that do not form cell-cell adherens junctions at all. In other words, we wanted to find out whether the formation of adherens (cell-cell) junctions between cells in an ensemble would influence their overall contractility. Experimentally disrupting the cell-cell border^22,28^ by calcium depletion or knock down β-catenin affect cell viability and introduce additional confounding factors. Therefore, we constructed a simple mechanical equivalent of a two-ASM cell ensemble with no cell-cell contacts. In order to do this, we considered isolated, single ASM cells cultured on the rectangular patterns as before. The traction fields for each single cell was first calculated independently. Then, we translated the coordinate system from the two independent traction force vector fields, such that the two ASM cells were adjoint and aligned in the same direction. The vector fields at matching grids were added up vectorially to create the new traction field of this theoretical ensemble. We then calculated the contractile moment of this theoretical two-cell pair (with no cell-cell junctions) and compared this to the contractile moment from our two-cell pairs at 300Pa and 13 kPa (Fig. 4). We found that at E=0.3kPa (healthy ECM, where the ASM-ASM cell coupling was maximum: see Fig. 3A and Fig. 2D), a connected two ASM cell ensemble has statistically significantly lower contractile strength (6.8±2.4 pNm, N =6) in comparison to two non-interacting ASM cells (17.8 ±8.9 pNm, N =10) p<0.05; Mann-Whitney Rank Sum Test, Fig. 4). This shows that ASM-ASM connections via adherens junctions are not merely a pathway for transmitting ASM force. Adherens junctions allow the contractile apparatus of individual cells to interact in such a way that the contractile strength of a multicellular ensemble is significantly lower than an equivalent ensemble which connects through cell-ECM focal adhesions rather than cell-cell adherens junctions. Comparing these results to the same scenario on E=13kPa (remodeled ECM, where the ASM-ASM coupling was significantly lower. see Figs. 3B&2D), the net contractility of a connected two ASM cell ensemble (15.9±5.9 pNm, N =4) was still lower than that of than two non-interacting ASM cells (21.1±10.5 pNm, N =10), but the difference was not statistically significant (p=0.127; t-test). As ASM-ASM coupling decreases, each ASM cell in the ensemble starts to behave like a single cell connected to a stiffer ECM and the net contractile moment increases significantly.

**Figure 4.**
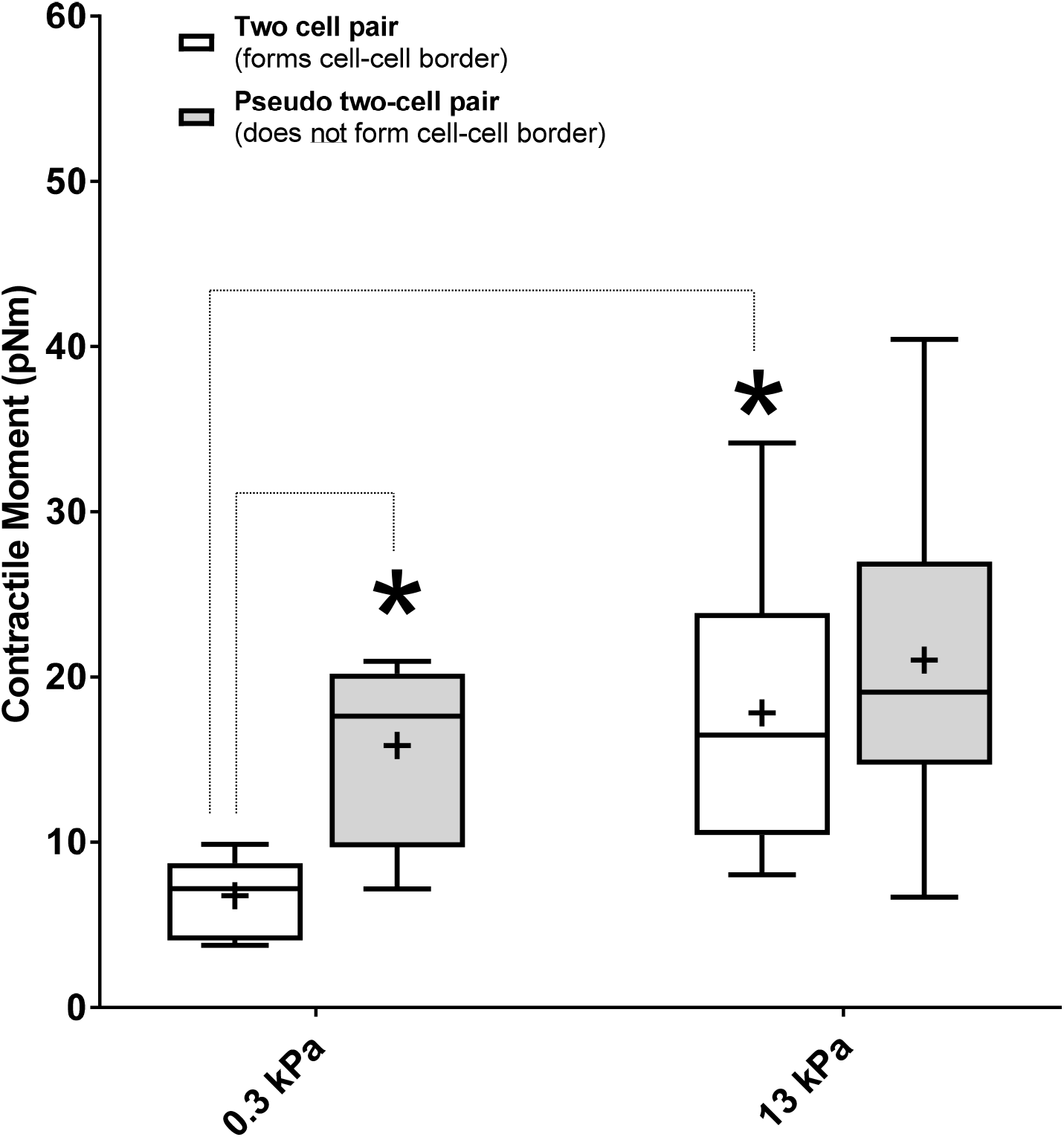
The contractile moment of a two-ASM cell ensemble is compared to that of a theoretical two-cell pair which does not form adherens (cell-cell) junctions. The ± sign indicates the mean value of each data set. At E =300Pa, (healthy ECM), the theoretical pair of two ASM cells with no cell-cell connections has statistically higher contractile moment than the two-cell pairs which form cell-cell borders (p<0.05, Mann-Whitney test). At E=13kPa the difference in contractile moment between the experimental two-cell pair and the theoretical pair is no longer statistically significant (t-test, p =0.379). The net contractile moment of the experimental two-cell ensemble does increase significantly with increase in ECM stiffness from E =300Pa to E =13kPa (Mann-Whitney test, p<0.05)

## Discussion

It is well understood by now that diseases such as hypertension, asthma and Chron’s are also accompanied by a substantial change in the extracellular matrix that surrounds, and connects to the smooth muscle cells^7^. The impact of such matrix remodeling on the excessive luminal narrowing of airways and blood vessels is not well understood. In this study, we measured the effect of ECM stiffness on the mechanical coupling between smooth muscle cells. We found that at low ECM stiffness mimicking healthy tissue (E=300Pa), ASM cells form cell-cell adherens junctions which create a well-defined border between the cells. Our force measurements showed that there is a strong mechanical coupling between the SM cells in the two-cell ensemble at this stiffness (Figs. 2D & 3A). These findings are inline with independent reports that the recruitment of β-catenin to N-cadherin is an integral part of the force transmission machinery in healthy airways^29^. As ECM stiffness was increased (E=13kPa), we found that there was a significant decrease in cell-cell coupling strength. Imaging confirmed a gradual loss of cell-cell adherens junctions in favor of cell-ECM focal adhesions with increased ECM stiffness. On very stiff ECM (E=40kPa), the cell-cell border completely disappeared and well-defined focal adhesions were observed. These measurements show that ECM remodeling can not only increase the force generated by the individual SM cells but also alter the connectivity among cells within the smooth muscle layer, with cells favoring a force transmission pathway that runs through the ECM.

We wanted to understand whether changes in connectivity among SM cells at the same ECM stiffness could alter the overall contractile properties of the multicellular SM ensemble. This is extremely difficult (if not impossible) to test experimentally without introducing additional confounding effects^22,28^. Hence, we created a virtual mimic of our two-cell ASM ensemble by repositioning two isolated single ASM cells to create an aligned dipole with no cell-cell contacts. Our results demonstrate that even at the same ECM stiffness, a change in connectivity among cells within the smooth muscle layer from cell-cell contacts to cell-ECM contacts would increase the overall contractile strength of the SM ensemble. While the method of using a theoritical two-cell pair is appropriate and accurate for the calculation of contractile moments, we do acknowledge the limitations that come with the theoretical nature of this calculation.

Our studies were limited to a two-cell ensemble system. In vivo, the airway SM cells are organized as a single layer of cells that spiral around the airways with a spiral pitch angle of 10 degrees ^19,30^. These cells are constantly subjected to periodic stretch associated with breathing. To what extent each these mechanical cues might impact our primary findings is therefore of great interest but is beyond the scope of the current work. Another area of future interest is to understand how SM cells respond to both contractile and relaxant stimuli. To this end, we are combining our approaches with concomitant measurements of pharmacological changes, to be performed in high-throughput, as is enabled by contractile force screening^20,31^

Here we used a novel UV based ECM patterning technique to create rectangular islands of gelatin on which we plated ASM cells to create our two-cell ensemble^32^. The technique was able to accurately reproduce a prescribed pattern on a PDMS substrate with a resolution of down to 2 microns. We also used a newly developed PDMS substrate, NuSil instead of polyacrylamide (PAA) gels which are commonly used in traction force microscopy measurements. NuSil has several advantages compared to PAA gels. NuSil allows for direct photopatterning of ECM proteins on its surface as opposed to an indirect transfer of protein using glass coverslips^33,34^. NuSil is also nonporous and optically clear (refractive index ~1.4). Most importantly, the stiffness of these substrates could be tuned within a range of Young’s modulus E=300 Pa-40kPa which allowed us to mimic the stiffness of healthy and remodeled human airway ECM stiffness. A detailed characterization of the mechanical properties of NuSil gels can be found in Yoshie et al. ^20^.

In this study, we found that increasing ECM stiffness from E=300Pa to 13kPa results in loss of cell-cell adhesions and weakening of the cell-cell coupling between airway smooth muscle cells. Comparing this result to previously published measurements in other cell types, the present findings appear to be cell-type dependent. Our results are consistent with disruption of bovine, canine and human endothelial cell monolayers due to substrate stiffening^35,36^, but not with measurements that show an increase in cell-cell coupling observed in myocardiocytes with an increase in matrix stiffness^17^. The cadherin-catenin complex/actin filament bonds that are at the core of cell-cell contacts have lifetimes that depend on force in a biphasic manner. The bond lifetimes increase with force up to 10pN (catch bonds/cell-cell adhesion is stabilized by force), after which the bond lifetimes decrease with increasing force (slip bonds/cell-cell adhesion is destabilized by force)^16^. This suggests that it is reasonable to expect the cell-cell coupling strength (Ψ) to depend on the contractility of cells considered, the range of ECM stiffness, the composition of the ECM and the specific focal adhesion & cell-cell adhesion proteins expressed in each cell type. We also note the complementary nature of cell-cell and cell-ECM contacts expressed by SM cells in an ensemble. A decrease in the area occupied by cadherin based cell-cell junctions was always accompanied by an increase in the total area occupied by focal adhesions. This observation is consistent with previous studies in other adherent cell types^37^.

In most existing models of luminal constriction^38,39^, the smooth muscle cells are lumped into one continuous layer in the cylindrical wall that generates the tension necessary for constricting the lumen while the extracellular matrix acts as the load against which the muscle constricts. In reality, these smooth muscle cells are organized in bundles and are able to dynamically adjust their connectivity with each other depending on extracellular mechanical factors such as matrix stiffness^19,30^. Here we have shown that the ECM stiffness can act as a switch that regulates whether forces are transmitted via the ECM or through cell-cell contacts. The change in connectivity can also significantly change the overall contractile strength of the ensemble. Excessive contraction of airways and blood vessels can therefore emerge as a result of change in connectiviity among SM cells driven by extracellular matrix remodeling. Our results highlight the need to develop new therapies for asthma and hypertension that target extracellular matrix remodeling.

## Methods

### Fabrication of NuSil Gel

NuSil (NuSil Silicone Technologies, Carpinteria, CA) parts A and B were mixed in a 1:1 ratio (by volume). An appropriate volume of part B crosslinker (V/V) from Sylgard 184 (Dow Corning, Midland, MI) kit was added to the solution to increase substrate stiffness to desired levels. The relationship between substrate stiffness and % of Sylgard 184 part B crosslinker to be added can be found in Yoshie 2018^20^. In our study 0.36% (V/V) of Sylgard 184 part B was added to create substrates with E=13kPa. Approximately 1mL of the mixture was placed on top of a 30mm coverslip and was spun coat onto the surface at 800RPM (Fig. 5A,B). Next, the coated coverslip was baked in an oven overnight at 60°C. Coverslips could then be coated with a thin layer of NuSil containing 1µm fluorescent beads for traction force microscopy (Fig 5C). The thin NuSil layer with beads was created by adding 1mL solution of 2µm fluorescent beads in a silicone solution (courtesy of Allen J. Ehrlicher, McGill University, Montreal, Canada) to the NuSil solution of a stiffness matching mixture. About 500µL of solution was added to the surface of the gel and spun coat at 2600RPM. The bead coated coverslip was again baked in an oven overnight at 60°C.

**Figure 5.**
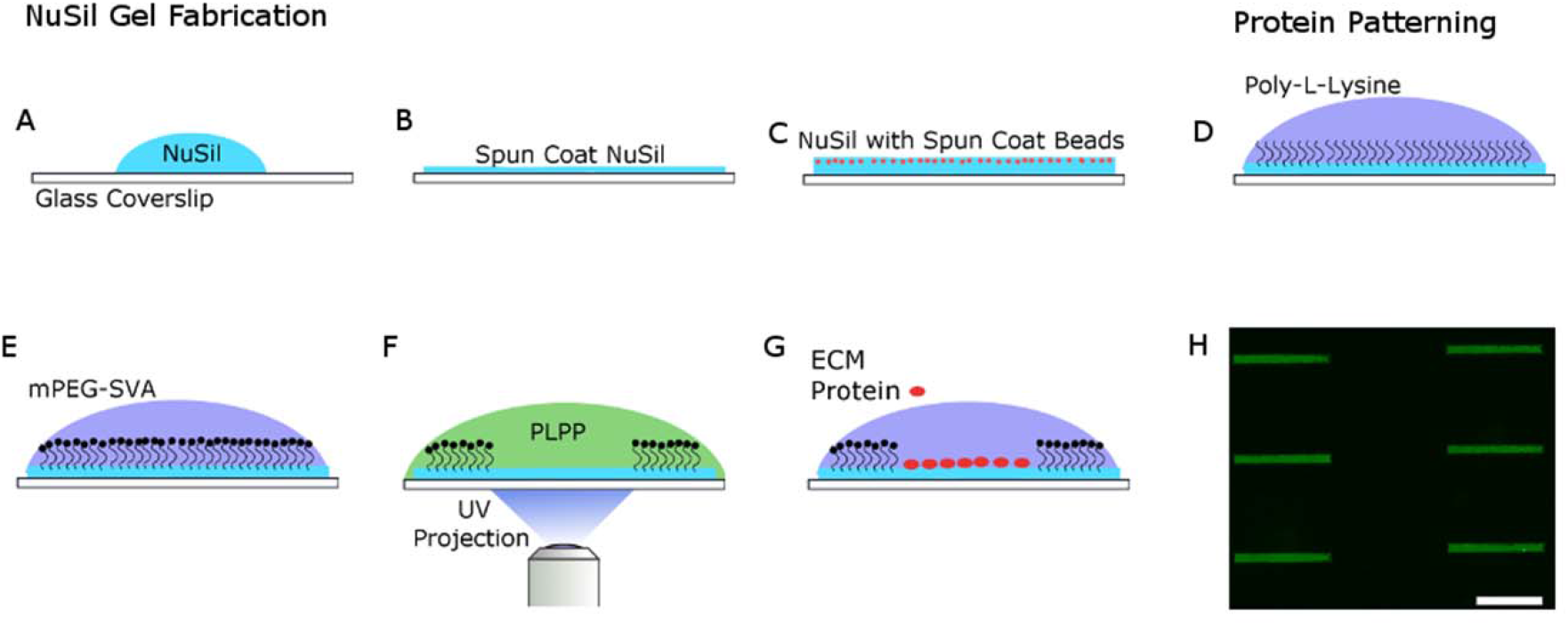
Schematic of the process used to create rectangular patterns of ECM proteins. **(A)**NuSil is spun coat onto a glass coverslip to create an even layer of gel on the surface of the cover slip as shown in panel **(B).** The NuSil is then baked overnight on an even surface at 60°C. The process can be repeated with the addition of fluorescent beads to create a uniform, thin layer of beads as shown in panel **(C)**. The surface is then coated with PLL (panel **D)** and then PEG-SVA (panel **E)**, which forms amide bonds to PLL, preventing cell adhesion outside the desired patterned area and to the PLL itself. **(F)** The desired pattern, a 15 µm x 150 µm rectangle, is then projected using UV light (λ=375nm) onto the surface of the NuSil gel in the presence of PLPP, which helps to rapidly degrade the PLL-PEG coating in the exposed area. **(G)** The desired ECM protein is then added to the surface of the gel and then rinsed to only adhere to the UV exposed area. **(H)** shows a representative image of 150x15 µm rectangular patterns of fluorescent gelatin. Scale bar is 100 µm.

### Photopatterning NuSil

NuSil gels were photopatterned using the Primo (Alveolé, Paris, France) micropatterning system. A schematic of the patterning process is illustrated in Fig. 5. The NuSil was coated with 0.1% Poly-L-Lysine (PLL; Sigma Aldrich, St. Louis, MO) for 1h at room temperature (Fig. 5D). After, the PLL was rinsed 3x with 1x PBS at pH7.2, followed by 3x of 10mM HEPES buffer at pH 8.2. Then, mPEG-SVA, MW 5kD (Laysan Bio; Arab, AL) was diluted to 50mg/mL in 10mM HEPES buffer at pH 8.2. The solution was added to the surface of gel and incubated for 1h (Fig. 5E). After incubation, the surface was rinsed with PBS. Then, PLPP at a concentration of 14.5 mg/mL (Alveolé) was added to the surface of the gel prior to patterning. The UV system was used to pattern 15 µm x 150 µm at a dosage of 1700 mW/mm^2^ (Fig. 5F) for traction force experiments. After patterning, the gel was then rinsed with 1xPBS and incubated with 0.1% gelatin (ECM protein used here) for 1h (Fig 5G). The samples were washed with 1x PBS, leaving gelatin only in the area patterned. A typical pattern obtained using this protocol is shown in Fig. 5H. Samples were then stored in the refrigerator until seeded.

### Cell Culture & Seeding Cells on Gel

Primary human airway smooth muscle cells (HASMCs) were acquired through the Gift of Hope Foundation (via Dr Julian Solway, University of Chicago). This is a publically available source and the donor remains anonymous and cannot be identified directly or indirectly through identifiers linked to the subject. This study meets NIH guidelines for protection of human subjects and no informed consent is necessary from this vendor as all donor identifiers are removed. All protocols used were carried out in accordance with the relevant guidelines and regulations that were reviewed and approved by the Institutional Biosafety Committee, Northeastern University. Cells were grown at 37 °C and 5% CO_2_and utilized prior to passage 7 for traction force experiments and for staining experiments. Cells were cultured in 10% fetal bovine serum, DMEM/F12, 1x penicillin/streptomycin, 1x MEM non-essential amino acid solution, and 250 µg/L Amphotericin B. Prior to measuring traction forces, the HASMCs were incubated in serum free medium for a minimum of 24 hours. The serum free medium was comprised of Ham’s F-12 media, 1x penicillin / streptomycin, 50µg/L Amphotericin B, 1x L-glutamine, 1.7mM CaCl_2_, 1x Insulin-Transferrin-Selenium Growth Supplement (Corning Life Sciences; Tewksbury, MA), and 12mM NaOH.

### Cell Traction Experiment

The protein coated gels were placed into 35 or 40mm interchangeable coverslip dishes (Bioptechs, Butler, PA). The gels were UV sterilized for 30 mins. Cells were seeded at a density of 2400 cells/well in 10% serum media for 15 mins. After 15 mins, non-adherent cells were removed by rinsing with 1x PBS and the media was replaced with serum free media. After 24h, cell traction forces were recorded by imaging the fluorescent beads using a Leica DMI8 microscope, a Leica DFC9000 camera and a Lumencore Sola SEII LED light source at 37°C with a 40x objective. Cells were then labeled with NucBlue (Fisher Scientific) live nuclear stain to determine the number of cells in the ensemble. After imaging, the cells were removed using RLT Lysis Buffer (Qiagen, Hilden, Germany). Cell traction forces were calculated using Fourier Traction Force Microscopy via a custom MATLAB (Mathworks, Natick, MA) software program^40^.

### Fluorescent Labeling of *β*-catenin and Vinculin

Cells were fluorescently labeled for vinculin (ab196454; Abcam, Cambridge, UK) and β-catenin (ab194119). Cells were fixed for 10min with 4% formaldehyde in PBS at room temperature. Cells were then permeabilized with 0.3% TRX-100 for 5min in PBS. Then, cells were blocked with 5% bovine serum albumen for 1h. They were then stained simultaneously for vinculin and β -catenin for 2h, both at 1:200 dilution ratios in 1XPBS.

### Statistical testing

Sigmastat (Systat Software, San Jose, CA) was used to perform statistical tests. All pairwise comparisons used the t-test when the data was normally distributed. Otherwise, the Mann-Whitney test was used to compare the median values. The specific tests used and the p-value are described along with the corresponding results. A p-value of 0.05 was used as the threshold for a statistically significant difference between data sets.

## Acknowledgements

This work was supported by NIH grants HL129468 and HL122513 (HP), and HL123552 (RK).

## Author contributions statement

HP, RK and SP contributed to the experimental design, SP & SS conducted the experiments, SS, SP, RK, and HP analysed the results. All authors contributed to manuscript preparation.

## Data availability

The datasets that support the findings of this study are available from the corresponding author on reasonable request.

